# Patient-specific computational models predict prognosis in B cell lymphoma by quantifying pro-proliferative and anti-apoptotic signatures from genetic sequencing data

**DOI:** 10.1101/2023.07.10.548371

**Authors:** Richard Norris, John Jones, Erika Mancini, Timothy Chevassut, Chris Pepper, Andrea Pepper, Simon Mitchell

**Affiliations:** Brighton and Sussex Medical School, Department of Clinical and Experimental Medicine, Brighton, United Kingdom; School of Life Sciences, University of Sussex, Brighton, United Kingdom

**Keywords:** Cancer, Computational Biology, DLBCL, Myeloma, Systems Biology

## Abstract

Genetic heterogeneity and co-occurring driver mutations impact clinical outcomes in blood cancers. Grouping tumours into clusters based on genetic alterations is prognostically informative. However, predicting the emergent effect of co-occurring mutations that impact multiple complex and interacting signalling networks remains challenging. Here, we used mathematical models to predict the impact of co-occurring mutations on cellular signalling and cell fates in diffuse large B cell lymphoma (DLBCL) and multiple myeloma (MM). Simulations predicted adverse impact on clinical prognosis when combinations of mutations induced both pro-proliferative and anti-apoptotic signalling. So, we established a pipeline to integrate patient-specific mutational profiles into personalised lymphoma models. Using this approach, we identified a subgroup (19%) of patients characterised by simultaneous upregulation of anti-apoptotic and pro-proliferative (AAPP) signalling. AAPP patients have dismal prognosis and can be identified within all current genomic and cell-of-origin classifications. Combining personalised molecular simulations with mutational clustering enabled stratification of patients into clinically informative prognostic categories: good (80% progression-free survival at 120 months), intermediate (median progression-free survival of 93 months), and poor (AAPP, median progression-free survival of 26 months). This study shows that personalised computational models enable identification of novel high-risk patient subgroups, providing a valuable tool for future risk-stratified clinical trials.

## Introduction

Mutational heterogeneity in haematological malignancies represents a major barrier to understanding the molecular mechanisms responsible for treatment response, and to the development of novel treatment strategies. With the advent of whole exome sequencing (WES) many malignancies, including B cell malignancies such as Diffuse Large B cell Lymphoma (DLBCL) and Multiple Myeloma (MM), have become characterised by interpatient mutational heterogeneity (1, 2). This heterogeneity contributes to variation in response to current treatments.

The most aggressive haematological malignancies frequently contain genetic aberrations affecting multiple genes, either through co-occurring mutations, e.g. double hit (DH) DLBCL or changes in the copy number of chromosomal regions containing multiple genes, e.g. gain1q multiple myeloma. DH DLBCL, featuring overexpression of *MYC* and *BCL2* (or *BCL6*), is among the most aggressive lymphoid malignancies, with very poor patient outcomes (3, 4). However, DH lymphoma represents fewer than 10% of DLBCL cases, while many more (30-40%) DLBCL patients relapse following frontline treatment (5). New approaches are needed to prospectively identify poor prognosis patients with the aim of developing more effective treatment strategies for this group.

Gene expression profiling split DLBCL into subgroups based on their putative cell of origin (germinal centre – GC, or activated B cell – ABC) (6). Subsequent studies leveraged WES to identify 5 or more prognostically informative patient clusters (7–9). These genetic groups have been arbitrarily named cluster 1 to 5 (C1-C5) or with names referencing the most mutated signalling pathways (MCD = MYD88 + CD79B), but are broadly consistent across multiple studies (e.g. C5 aligns with MCD) (7–9). Clustering patients in this way largely ignores the complexity of the molecular signalling networks that may be impacted by their individual genetic landscapes. For example, a patient with most mutations converging on NF-κB is likely to be assigned to C5 (7). However, this assignment ignores the presence or absence of functionally significant co-occurring mutations in other pathways that may impact prognosis.

Computational models of molecular signalling in normal B cells have been used to predict cell proliferation and survival; predictions that have been validated by in vitro laboratory experiments with single cell resolution (10–13). Furthermore, incorporating mutations, and the impact of mutations on protein abundance/activity, into these models predicts cellular responses in experimental assays (10, 11, 14, 15). However, it is not known whether these models can predict outcomes at the individual patient scale, nor whether mutational data alone is sufficient to enable *in silico* simulations to make clinically relevant predictions in blood cancers. We hypothesise that contextualising sequencing data within patient-specific, virtual signalling networks may more precisely delineate the consequent cell fate decisions that impact prognosis.

In this study we used mechanistic computational models to simulate how mutations combine in B cell malignancies, and determined whether these approaches could predict patient outcome. We developed a pipeline to create individual patient simulations using WES data, and tested whether these personalised models could generate clinically informative prognostic information.

## Materials and Methods

Detailed computational methods, and a lay summary of the methods to generate all computational figures, are provided in the Supplementary Material. Methods are summarised below. Code (as Jupyter notebooks) used to generate and run all computational models, including descriptions of the reactions, parameters, and rate laws of each model, and code to plot output are available in the Github repository (https://github.com/SiFTW/norrisEtAl).

### Model generation

We employed established, computational models of healthy B cells, which enable simulation of proliferation, apoptosis and terminal B cell differentiation in a heterogeneous B cell population (10, 11, 13). The models consisted of a series of differential equations, representing the rate of change of biomolecules (mRNAs, proteins, or protein complexes) over time. Models were converted to Julia to be solved using DifferentialEquations.jl (16, 17).

### Cell cycle and apoptosis modelling

The molecular networks representing apoptosis and the cell cycle were isolated from the comprehensive B cell model (10, 11, 18). Cell-to-cell variability was simulated by distributing model parameters as described previously (10). Over-expression of cMyc and Bcl2 was simulated by increasing the parameter controlling the transcription rate of the respective mRNA. An equilibrium phase was simulated prior to all time course simulations. For the cell cycle model, the equilibrium phase ended when the same cell mass was reached prior to each cell division for 5 cycle (within 3 decimal places). The final state of the equilibrium phase was used as the initial condition for the time course phase. Cell death was defined as the first time point at which over 10% of PARP was cleaved.

### Multi-scale model

The NF-κB component of the established B cell model was updated to include reactions from a more recent and comprehensive model of NF-κB (12). Apoptosis and cell division were triggered as published previously (cleaved PARP >2500 triggers apoptosis, CDH1 >0.1 triggers cell division) (10, 11). Previous studies introduced two separately simulated daughter cells with each cell division (10, 11, 18), however here this resulted in an unfeasible simulation size in highly proliferative simulations. Cell fates have been shown to be reliably inherited in B cells such that daughter cells will achieve similar fates at similar times (16). Therefore, at each cell division we replaced the mother cell with a single simulation representing 2^(generation-1)^ daughter cells (2 cells following first division, 4 cells following second division etc.).

### Modelling patients

Mutation profiles identified from WES and clinical data were downloaded through the cBioPortal (7). We restricted the analysis to genetic changes impacting genes that could be mapped to modelled parameters (https://github.com/SiFTW/norrisEtAl/blob/main/geneList.txt) and leveraged OncoKB to simulate the impact of mutations annotated as “likely oncogenic” (31). We created 113 patient-specific models (data for 135 patients available, 24 patients did not harbour any mutations that mapped to model parameters). We assumed half of the normal expression rate of each mRNA could be attributed to one of the two copies of the gene. Therefore, parameters halved for copy number loss and loss-of-function mutations and increased by half (1.5-fold change) for copy number gain and gain-of-function mutations. Copy number alterations of chromosomal regions were modelled by identifying each gene that could be mapped to model parameters within the chromosomal region. Multiple mutations affecting the same parameter were combined multiplicatively, for example a patient with a MYD88 mutation and a CD79B mutation (both increasing NEMO:IKK activity by 1.5 fold individually) was simulated as 2.25-fold increased NEMO:IKK activity. The effect of mutations in genes not modelled directly was assigned to the closest modelled molecular species: e.g. *MCL1* is modelled as overexpression of *BCL2*, and *CARD11* mutations are modelled as increasing NEMO-IKK activity. As biomolecules such as BCL2/cMYC mRNA represented the sum effect of multiple mRNAs, e.g. MCL1+BCL2, these were renamed anti-apoptotic/pro-proliferative (AAPP) factors respectively, to avoid inaccurate comparisons with experimental measurements of these transcripts. Patient simulations were performed for 12 simulated hours, on a single cell with all parameters consistent across all patients other than those impacted by mutations. The concentration of pro-/anti-apoptotic factors at 6h was used for patient stratification, with patients stratified as above or below the mean concentration at that time point.

### Downstream analysis

All model outputs were analysed, and plots generated using the Julia programming language. Scripts relating to each figure. can be found in the corresponding folder available in the GitHub repository (https://github.com/SiFTW/norrisEtAl/).

## Results

### Simulating individual mutations recapitulates experimental measurements

To determine whether established computational models of B cell molecular signalling could make predictions of patient outcomes we first focused on DH DLBCL. The apoptotic regulatory network was isolated from an established B cell model, and we simulated the impact of *BCL2* overexpression on apoptotic signalling (11, 18)(Fig. 1A). A gene dose-dependent delay of apoptosis measured by poly ADP-ribose polymerase (PARP) cleavage up to the equivalent of 5 extra copies of *BCL2* was seen in these simulations, beyond which the cells survive the full 140 hours of the simulation (Fig 1B). We repeated these simulations in a population of 1000 molecularly heterogeneous individual B cells (Fig. 1C and S1A). In these simulations cell death times retained a log-normal distribution, but that 1.5-fold *BCL2* over-expression increased the mean survival time in individual B cells by 1.3-fold (Fig. 1C). This is consistent with the delayed cell death seen in B cells from Bcl-2-overexpressing mice (19).

**Fig. 1.**
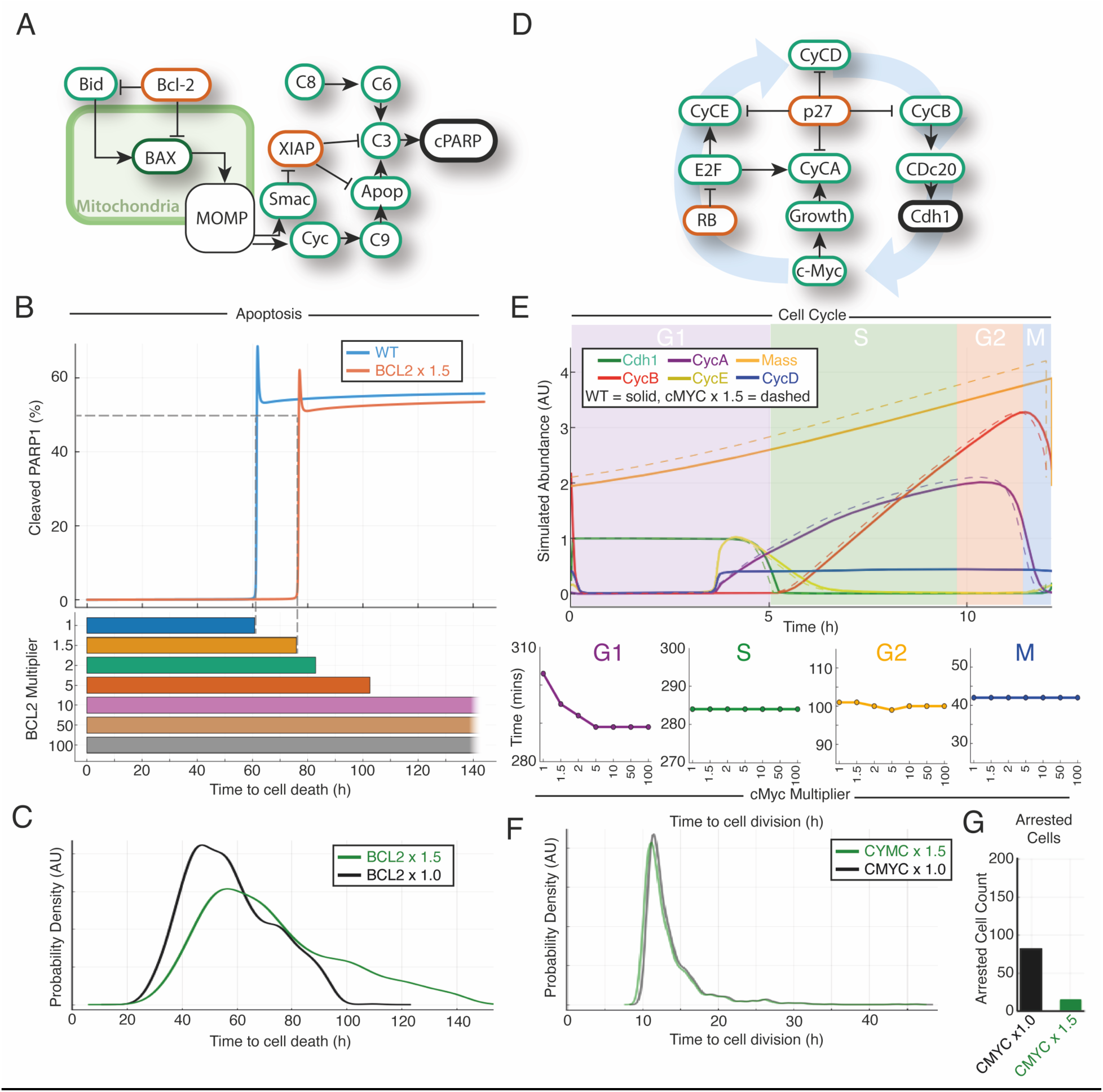
Computational modelling of the cell cycle and apoptosis reveal limited impact of archetypal double hit mutations on their respective molecular networks. **A-C** Apoptosis model simulations. **D-F** Cell cycle model simulations. **A** Schematic of the apoptosis model leading to cleavage of PARP. **B** (top) The percentage of cleaved PARP over time for two simulations (WT and BCL2 overexpressed) using the apoptosis model. There is a delay in PARP cleavage in the presence of overexpressed BCL2. **B** (bottom) Increasing time to death for cells with increasing BCL2 expression. **C** Graph showing distribution of time to death in simulation of 1000 cells, unmutated (black) compared to 1.5 fold BCL2 upregulation (green). **D** Schematic of the impact of cMyc on the cell cycle model. **E** (top) Output of the abundancies of different cell cycle proteins run to a limit cycle for a WT cell (solid line) and a cell in which cMYC is overexpressed (dotted line) showing a slight shortening of the cell cycle for some. CycA/D/E = Cyclin A/D/E, Cdh1= cdc20 homolog 1. E (bottom) Time for completion of cell cycle phases for cells with varying cMYC expression showing the main effect is in G1. **F** Distribution of time to cell division for simulations of heterogeneous populations of 1000 cells. Unmutated (black) compared to 1.5 fold cMYC upregulation (green) show very little difference. G Number of cells, from the simulations in panel **F**, for which the cell cycle has arrested in unmutated cells and cells in which cMYC is upregulated 1.5 fold. Upregulated cMyc substantially reduces the number of cells in cell cycle arrest.

*cMYC* is a proto-oncogene that plays a role in tumour cell growth and has previously been modelled as acting on B cell progression within the cell cycle (13, 20). Simulating *cMYC* in this way has been found to recapitulate both single-cell and bulk experimental measurements (10, 11). We isolated the cell cycle regulatory network from an established model of B cells and simulated the impact of overexpression of *cMYC* (11, 18) (Fig. 1D). These simulations predicted that increasing *cMYC* expression by the equivalent of 1 extra copy (1.5-fold increase) shortens the cell cycle by just 8 minutes in a 12-hour cycle; an effect size much smaller than expected (Fig. 1E). The sensitivity analysis also found that *cMYC* expression had a smaller effect on cell cycle time than other genes in the pathway (Fig. S2). Importantly, this result recapitulated experimental tracking of division times in B cells transfected with *cMYC* and cellular proliferation measurement in lymphocytes from Eμ-Myc mice (21). In simulations of increasing levels of *cMYC* expression, the small decrease in total cell cycle time was primarily due to a shortening of G1 phase, with a gene dose-dependent effect up to the equivalent of 5-fold higher expression, after which no further effect was seen (Fig. 1E). Upregulating *cMYC (x1.5)* in a heterogeneous population of 1000 individual cells confirmed these findings and resulted in simulated cell cycle timings that recapitulated the distribution of cell cycle duration measured by time-lapse microscopy in murine B lymphocytes (22) (Fig. 1F and Fig. S1B). In these simulations, increased *cMYC* expression was found to have a minor effect in all cells, decreasing total average cell cycle time by just 11.5 minutes out of a 12-hour cycle (Fig 1F).

The wild-type simulation identified a population of cells (8%) in cell cycle arrest. Concentrations of the cell cycle inhibitor p27, and complexes containing p27, were upregulated in these arrested cells (Fig. S3), which recapitulates experimental measurements of cell cycle arrest (24). Interestingly, in simulations of 1.5-fold *cMYC* overexpression the cell cycle arrest was reversed in over 80% of these cells (Figure 1G). The number of cells rescued from arrest by *cMYC* overexpression was proportional to the level of *cMYC* expression (Fig. S4) and these rescued cells became the most rapidly proliferating cells within the cell population (Fig. 1G and S1B). In keeping with these findings, *cMYC* is amongst the most differentially expressed genes in quiescence and reverses quiescence in haematopoietic stem cells (23, 24). Recent data has demonstrated that increasing the abundance of cMyc can enable senescent cancer cells to resume division (25). Taken together this data shows that computational modelling of the impact of *cMYC* and *BCL2* mutations, on the cell cycle and apoptosis respectively, accurately recapitulates multiple experimental measurements.

### Multi-scale modelling predicts that mutations that converge on apoptosis and the cell cycle confer poor prognosis in blood cancer patients

To establish whether computational simulations can accurately predict patient outcomes, we initially simulated the well-characterised subgroup of lymphoma patients with mutations affecting *cMYC* and *BCL2* (DH lymphoma). An agent-based multiscale model, previously used to simulate immune responses, was used here (10, 11, 13). In this framework NF-κB-activation through NEMO-IKK provided the input, resulting in the induction of NF-κB target genes (e.g. CYCD, cMYC, IRF4, BCL2). The impact of these target genes on the molecular networks controlling cell division and death could trigger the cell cycle to complete mitosis (resulting in daughter cells being added) or the cell to undergo apoptosis (resulting in the cell being removed, Fig. 2A and S5, see methods). Given that normally two copies of each gene are present we assumed half (0.5-fold) the gene expression could be attributed to each copy. Simulations with upregulated (equivalent to 1 extra copy, 1.5-fold increased expression) *cMYC* and *BCL2*, resulted in cell number increases due to increased cell division and survival respectively. When both genes were upregulated (1.5-fold expression) cell counts increased in an additive manner (Fig. 2B, left). However, a 4-6 fold increase in expression of *BCL2* and c*MYC* is commonly observed in these genes when they are mutated in lymphoma patient samples (26). Under these conditions (5-fold increased expression), the simulations resulted in an exponential increase in cell numbers (Fig. 2B, right). The classification of DH lymphoma includes mutations in *cMYC* and *BCL2* or *BCL6*. Performing the same simulation on the *cMYC* + *BCL6* DH lymphoma resulted in a substantially smaller effect compared to the *cMYC* + *BCL2* DH (Fig. 2C). Using cell numbers as a proxy for disease prognosis would result in predictions that *cMYC* + *BCL2* DH lymphoma would have worse prognosis than *cMYC* + *BCL6* DH lymphoma (Fig. 2B and C). A multicentre retrospective data collection study, published while this study was being conducted, confirmed our model prediction by showing that *cMYC* + *BCL6* DH lymphoma does not represent high risk lymphoma, while *cMYC* + *BCL2* DH lymphoma has particularly poor prognosis (Fig. 2D).

**Fig. 2.**
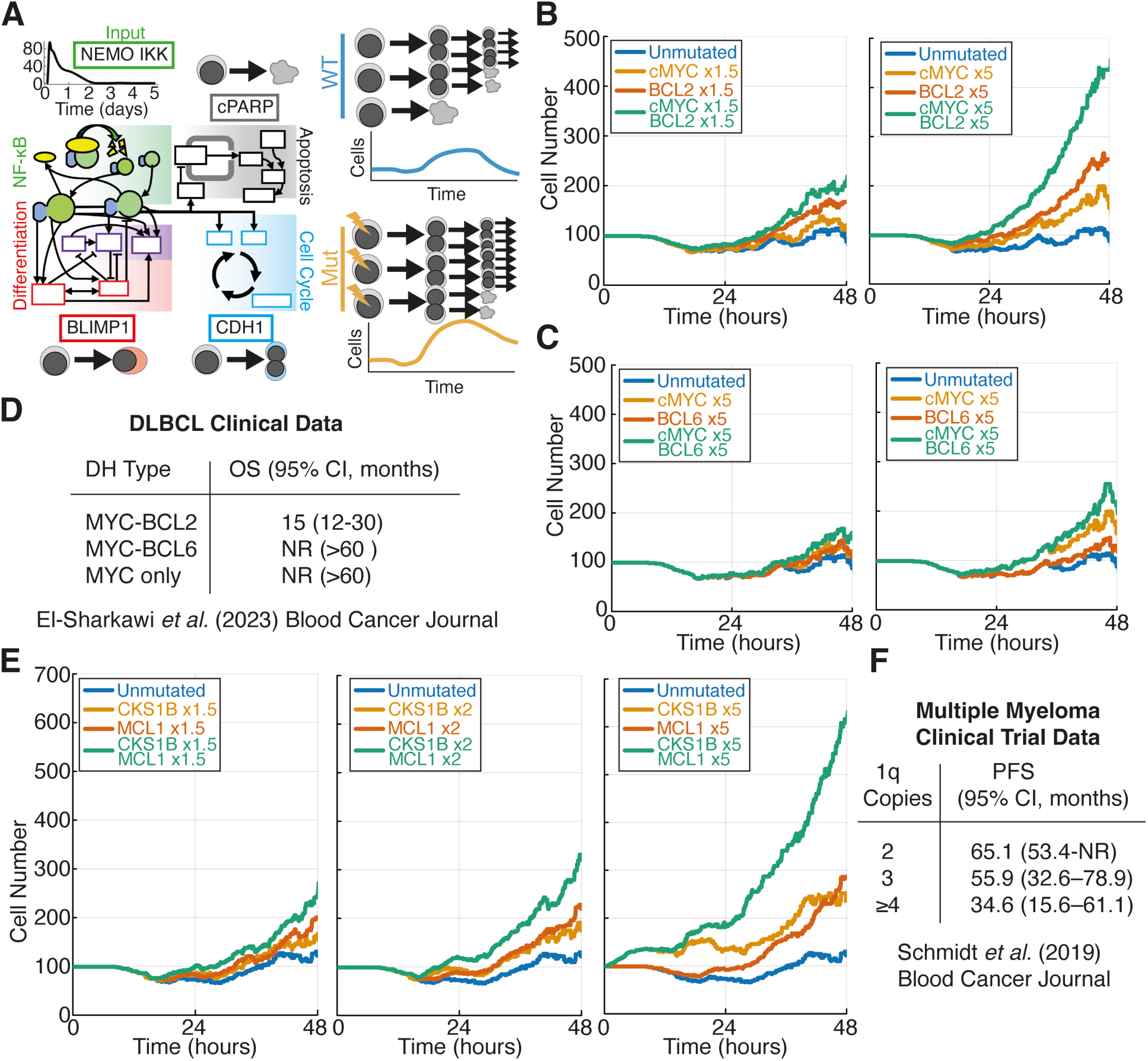
Multi-scale modelling of mutations in subsets of DLBCL and Multiple Myeloma recapitulates clinical trial data. **A (left)** Simplified schematic model divided into component signalling pathways featuring NFκB signalling, apoptosis, the cell cycle and differentiation (adapted from ref 11), full detail provided in Supplementary Material. **A (right)** Schematic representation of the progression of cell lineages over time during multiscale simulations. The example depicts a mutation that allows cells to continue to proliferate when they would otherwise die. **B&C** Simulated cell population size (cell count) over time for wild type (blue), individual mutations (orange and red) and double hits where both mutations are combined (green). **D** Overall survival (OS) data for groups of patients with MYC and BCL2/BCL6 mutations. **E** Cell population size (cell count) over time for simulations of gain1q multiple myeloma. CKS1B and MCL1 are located within chromosome 1q and therefore amplifications in this chromosome increase the abundance of both of these genes. **F** Progression-free survival (PFS) data for groups of patients with upregulation of CKS1B and MCL1 due to 2, 3 or 4 copies of chromosome 1q. NR = not reached.

To determine whether computational predictions could recapitulate disease prognosis in other high-risk haematological malignancies we simulated gain 1q multiple myeloma (Fig. 2E). Gain 1q MM is a commonly occurring cytogenetic event in MM associated with therapeutic failure and inferior prognosis (27). Genes encoding the Bcl2-family protein Mcl1 and cell cycle regulator Cks1b both reside on chromosome 1q21 and have been implicated in the pathogenesis of gain1q MM (28). Simulating the upregulation of each gene individually and in combination revealed a dose-dependent increase in cell numbers as the number of copies of genes on 1q increased (Fig. 2E). Retrospective analysis of clinical data found a dose-dependent worsening prognosis with increasing copies of chromosome 1q (29) (Fig. 2F). Strikingly, with a 5-fold increase in expression of *MCL1* and *CKS1B* the cell population continues to increase beyond the presence of any input signal (IKK activity returns to basal by 72 hours, Fig. S6). These simulations indicated that amplification of the chromosomal region containing *MCL1* and *CKS1B* may result in constitutive signalling and microenvironmental independence, which is circumstantially supported by the observation that 1q21 is amplified in 91% of MM cell lines (30).

We noted that within the models we were using, *cMYC* and *BCL2* in DH lymphoma impact the cell cycle and apoptotic regulatory networks respectively, while *BCL6* is situated within molecular networks controlling terminal B cell differentiation (Fig. S5). In the context of gain1q MM, *CKS1B* and *MCL1* over expression results in the perturbation of the cell cycle and apoptosis respectively. Consistent with DH lymphoma, we found that when mutations simultaneously impact the cell cycle and apoptosis they combined deleteriously in simulations and conferred poor prognosis in clinical data.

### A computational pipeline enables simulations of heterogeneous patients from WES data by mapping mutations to simulation parameters

Having demonstrated the model can accurately predict known poor prognosis mutation combinations, we next sought to establish whether other co-occurring deleterious mutations that impact both the cell cycle and apoptosis could be identified through computational modelling. Comparing simulations of activating mutations in *BCL2* and *cMYC* to simulations of an upstream NEMO:IKK/NF-κB–activating mutation demonstrated that NF-κB-activating mutations could cause an increase in both BCL2 and cMYC mRNA similar to that seen in double hit mutation lymphomas (Fig. 3A). To employ the model to identify additional combinations of mutations that could result in simultaneous anti-apoptotic and pro-proliferative molecular signalling, we sought to leverage a WES dataset that captures the mutational heterogeneity in individual DLBCL patients (7). We developed a pipeline to map both individual gene-level mutations and chromosomal arm-level copy number alterations to computational parameters in the multi-scale model (Fig. 3B, see Materials and Methods). This process created 113 patient-specific models, with parameters modified based on gene alterations described from analysis of WES data (Fig. 3B), while all other parameters remained the same (31).

**Fig. 3.**
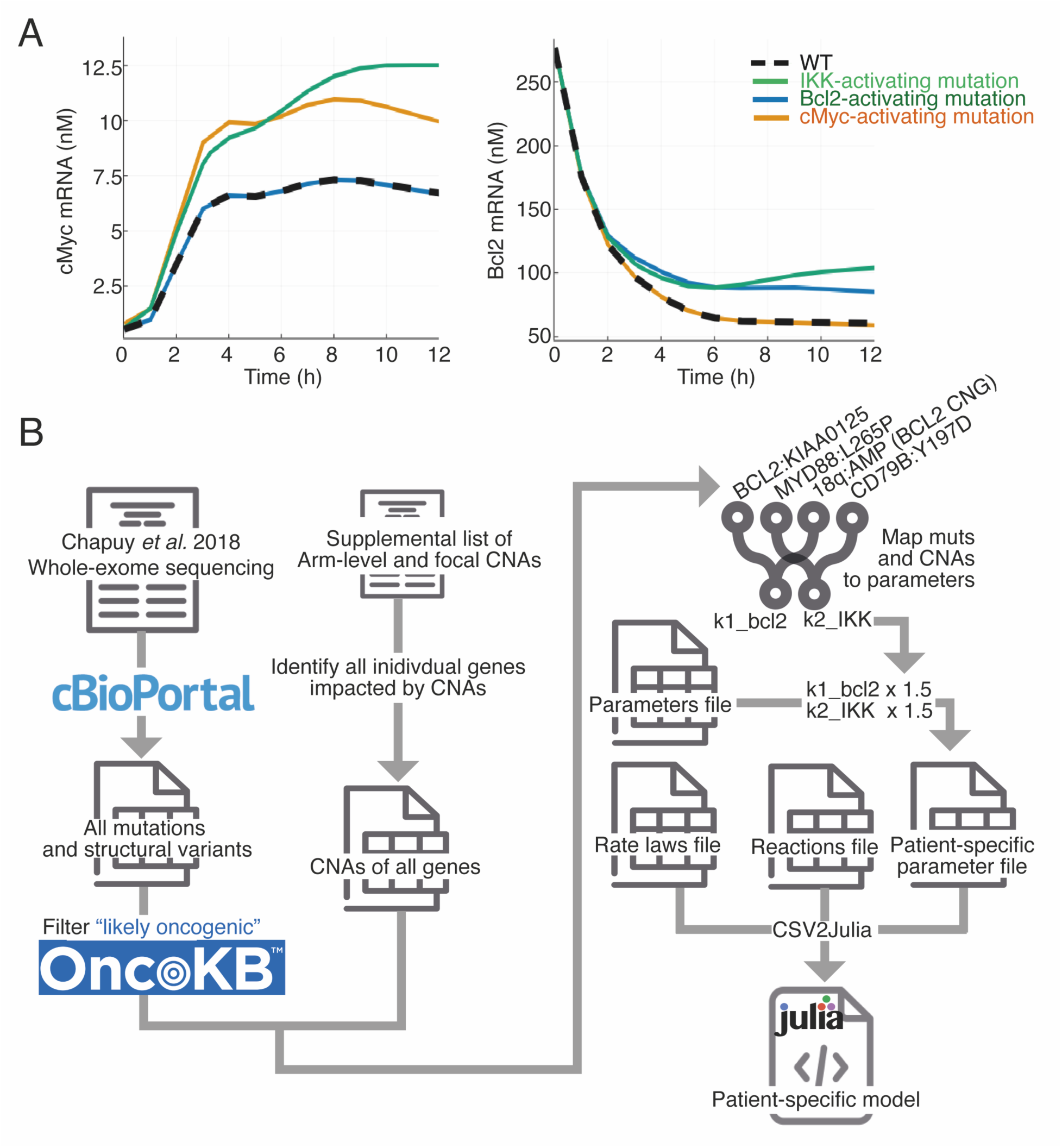
Multiple mutations can create anti-apoptotic and pro-proliferative signalling. **A** Abundance of cMyc (A) and Bcl2 (B) mRNA in simulations of WT (dash), an IKK-activating mutation (green), a Bcl2-activating mutation (blue) and a cMyc-activating mutation (orange) over time. **B** Pipeline to incorporate mutational events from genetic sequencing to create patient specific models. Example mutational mappings are shown including the model parameters they modify. The full mapping is provided on in the Github repository (https://github.com/SiFTW/norrisEtAl/blob/main/muts2Params.csv).

### Personalised patient simulations identify an anti-apoptotic and pro-proliferative subgroup of patients with poor prognosis

Analysing pro-proliferative and anti-apoptotic factors over time for individual patient simulations revealed highly heterogeneous expression between patients, with most dynamic changes occurring in the first 6 hours (Fig. 4A). Concentrations at 6 hours were chosen to stratify patients as this was the earliest time point at which most transient dynamics had passed but cells had not been removed from the simulation due to cell division or cell death. Furthermore, concentrations at later time points correlated strongly with those at 6 hours (R^2^ > 0.97 between 6h, 9h and 12h time-points for both pro-proliferative and anti-apoptotic factors, Fig. S7). Patients with above or below mean average expression of pro-proliferative or anti-apoptotic factors alone at 6 hours did not differ in their progression-free survival (Fig. 4B, p>0.05). But stratification of patients based on the combination of these factors identified a group of patients (19% of the cohort) for which both anti-apoptotic and pro-proliferative (AAPP) factors were upregulated (Fig. 4C, example shown in orange in Fig. 4A). Patients classified as AAPP using computational simulations had significantly worse prognosis than DLCBL patients not classified as AAPP (Fig. 3D, median PFS 26 months compared to not reached, p=0.001). The hazard ratio for AAPP patients was 2.767 (95% confidence interval: 1.259-6.080) when compared to patients not assigned AAPP (95% confidence interval: 0.664-1.506)

**Fig. 4.**
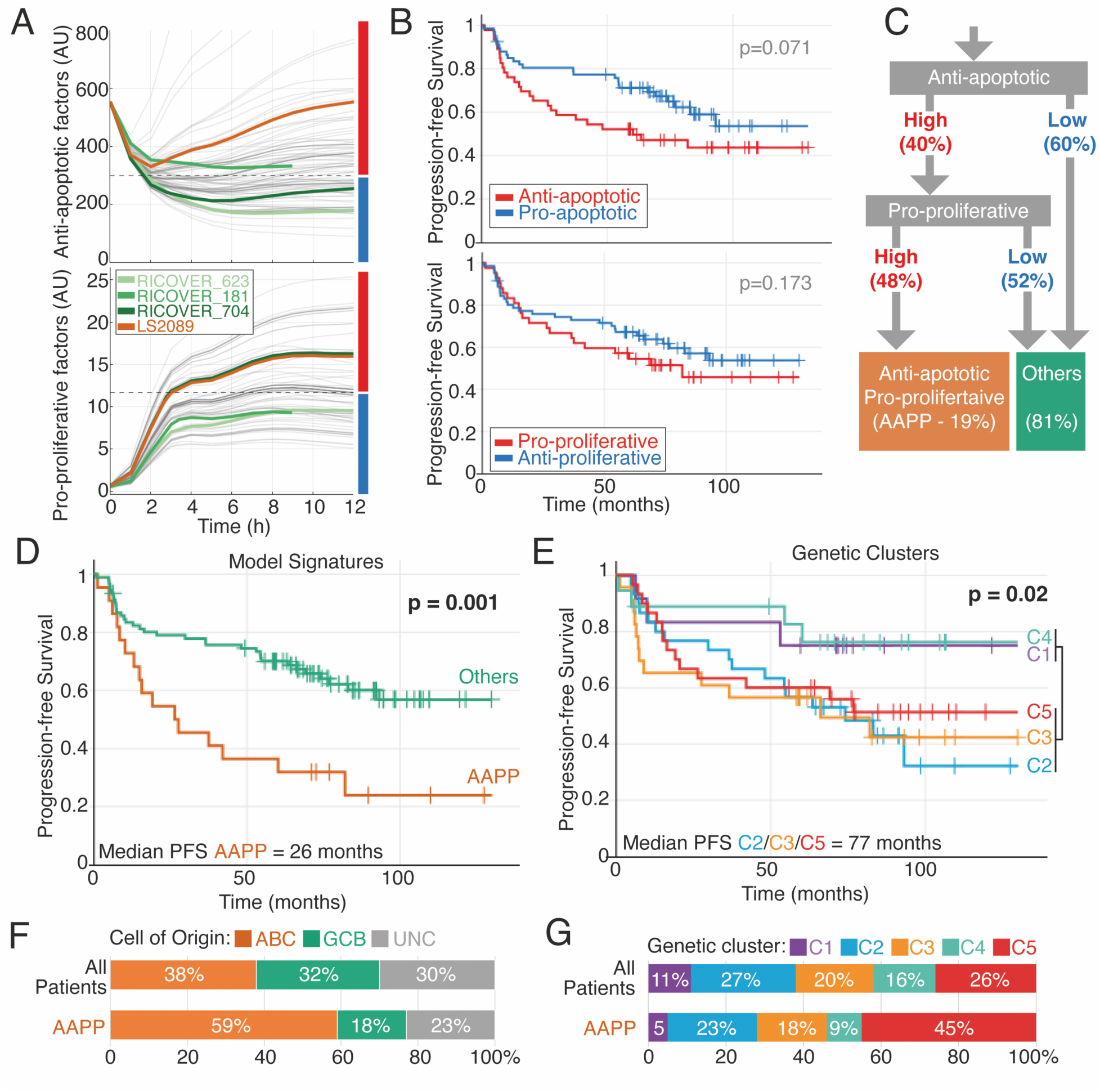
Patient-specific modelling and stratification by pro-proliferation and anti-apoptotic species identifies poor prognosis patients within all cell-of-origin and genetic clusters. **A** Abundance of anti-apoptotic and pro-proliferative factors for individual patients (grey) simulated using the pipline in Fig 3. Example patients, labelled with their anonymised identifier, in each of 4 groups defined by the relative abundance of these factors are highlighted: low/low = light green; high anti-apoptotic = mid green; high pro-proliferative = dark green; anti-apoptotic and pro-proliferative (AAPP) = orange. The dashed line indicates the mean value at 6h and is used for stratification into high and low in subsequent panels. **B** Kaplan-Meier (KM) plots comparing progression-free survival for DLBCL patients with model-identified high or low anti-apoptotic factors (top) and high or low proliferative factors (bottom). **C** Flow chart describing the identification of anti-apoptotic pro-proliferative (AAPP) patients. **D** KM plot for AAPP patients and all other patients. **E** KM plot for all modelled DLBCL patients grouped according to mutational clusters from ref. 7. **F&G** Grouped bar plots showing the percentage of patients in the AAPP group vs all patients, stratified by cell of origin **(F)** and genetic clusters **(G)**. ABC = Acitvated B Cell, GCB = Germinal Center B cell, UNC = unclassified.

Comparing stratification through computational modelling with clustering of WES data revealed that computational modelling enabled the identification of a substantially worse prognosis patient cluster (AAPP) than could be identified from mutations alone. The AAPP cluster had a median PFS of just 26 months compared to 77 months in the poor prognosis clusters (clusters 2/3/5 = C2/C3/C5) identified in reference 7 (Fig. 4D and E). Comparing cell of origin (COO) and genetic clustering in the entire patient cohort with AAPP patients showed that AAPP patients spanned all COO classifications and genetic clusters, but AAPP patients were enriched in poor prognosis COO (ABC) and genetic cluster 5 (Fig. 4F and G).

Importantly, AAPP patients identified by modelling could not be identified from mutational burden alone without simulation, and did not differ in their age at diagnosis, IPI, CNS involvement, mutation count, number of driver mutations, number of copy number alterations, or ploidy (q=0.1-1.0 for all clinical metrics) compared to the rest of the cohort (Fig. S8).

Only one mutation, Myd88:L265P, was significantly enriched in the AAPP patients (false discovery correction using Benjamini-Hochberg, q=0.00027). The computational modelling had not simply re-identified MYD88:L265P patients as the presence of Myd88:L265P was not sufficient to confer poor prognosis in this cohort (Fig S9A). However, we did observe bi-modal progression free survival among MYD88:L265P patients (Fig S9A and B). Strikingly, among MYD88:L265P patients, those identified as AAPP had significantly worse PFS (Median PFS MYD88:L265P+AAPP = 13.08 months, MYD88:L265P+not_AAPP=NR, p=0.0081), and overall survival (median OS PFS MYD88:L265P+AAPP = 26.8 months, MYD88:L265P+not_AAPP=NR, p=0.0081)(Fig S9C and D). AAPP patients are a larger subgroup (19%) than patients with either DH or triple hit lymphoma (<5%)(32). Surprisingly, computational modelling did not assign patients with concurrent structural variants affecting MYC and BCL2 to the AAPP category. However, the cohort only contained 3 such patients was therefore not powered to assess their outcome. Furthermore, it is worth noting that all three patients had low IPI (≤1) and none of these patients had progressed at the end of the study. This suggests that computational modelling of patients can identify poor prognosis patients independent of double hit structural variant status.

### Combining personalised computational simulations with consensus clustering enables prognostic stratification

When comparing mutational clustering with model-based stratification we observed that the model had greater power to identify poor prognosis patients while mutational clustering excelled in identifying good-prognosis patients (C1 and C4, Fig. 4E, had substantially better outcomes than non-AAPP patients Fig. 4D, “Others”). Therefore, we devised a patient-stratification approach using results from both methods, ultimately assigning patients to 3 patient categories (AAPP=19%, C2/C3/C5 not AAPP = 57%, C1/C4 not AAPP = 24%, Fig. 5A and B). PFS in these three clusters was significantly different (p=0.001), with AAPP patients having dismal prognosis (PFS 26 months), C2/C3/C5 not AAPP patients having intermediate prognosis (PFS 93 months), and C1/C4 not AAPP patients having good prognosis (with 10-year PFS > 80%) (Fig. 5C). This stratification approach was also robust to the inclusion or exclusion of patients that did not contain a single mutation that mapped to a model parameter (Fig. S10).

**Fig. 5.**
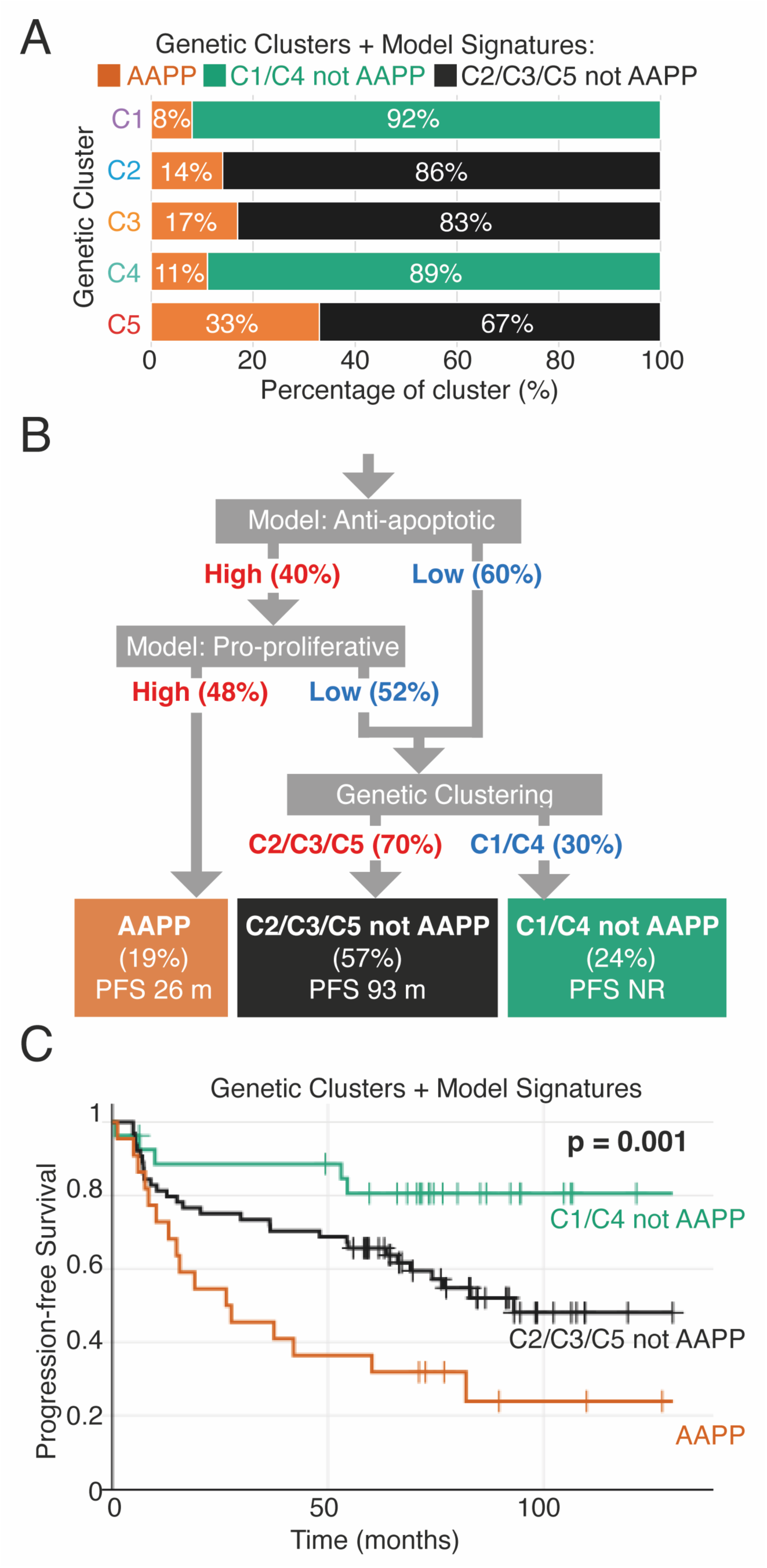
Combining patient-specific modelling with genetic clustering stratifies patients into distinct prognostic clusters. **A** Grouped bar plot showing the percentage of AAPP and non-AAPP patients in each genetic cluster from ref. 7. **B** Flow chart describing stratification of patients first using patient-specific models to identify AAPP patients and then using clustering to stratify non-AAPP patients. **C** Kaplan-Meier plot of patients stratified using flow chart in panel B.

## Discussion

Predicting which blood cancer patients will respond well to treatment is key to empowering patients and clinicians in clinical decision making and may be required to design trials that advance the standard of care in heterogeneous blood cancers. We found that leveraging genomic sequencing data from DLBCL patients to create personalised simulations and combining this analysis with mutational clustering could outperform either approach in isolation. Molecular clustering of DLBCL by multiple groups has identified clusters of patients that differ in the signalling pathways most impacted by mutations (7–9). A single mutation may impact multiple signalling networks, and multiple mutations may overcome or exacerbate each other. For example, gain of 19q increases BAX (pro-apoptotic) and CCNE1 (pro-proliferative), NFKBIB (pro-apoptotic and anti-proliferative). Computational modelling enabled identification of patients with co-occuring mutations that are simultaneously pro-proliferative and anti-apoptotic, which could not be determined from mutational clustering alone. Importantly, this modelling approach identified patients with worse prognosis (PFS of 26 months) than genetic clustering and even artificial intelligence-based approaches informed by more data (gene expression data, mutational data, age and sex, poor prognosis cluster PFS 43.23 months)(32). We believe this highlights the importance of encoding molecular network knowledge to contextualise mutational information (12, 33).

There is compelling evidence supporting the adoption of genetic sequencing at diagnosis of DLBCL to determine prognosis (34, 35). All mutations and CNAs modelled in this study affected recurrently mutated genes, which are profiled by targeted sequencing panels. As such, the data required for modelling could be generated at low cost and from fixed diagnostic samples, and potentially even a plasma sample (36). Furthermore, the recent development of alternatives to R-CHOP is likely to motivate utilisation of genomic sequencing data to rationally assign therapies. There is emerging evidence supporting the addition of polatuzumab vedotin for high risk DLCBL, the addition of Bortezomib in ABC-DLBCL, or the use of dose-adjusted EPOCH plus rituximab for patients expressing high levels of MYC and BCL2 (37–39). Factoring in the decreasing costs of sequencing, it seems likely that molecular profiling will become standard diagnostic practice in DLBCL, with the aim of precisely identifying patients that would benefit from an alternative to R-CHOP. Here we show that computational modelling can improve the ability to identify such patients without requiring additional data.

More work is required if modelling is to become a widely used tool for personalised medicine approaches. While computational simulations identified poor prognosis patients (AAPP), mutational clustering was still needed to identify good-prognosis patients (C1/C4). Here we chose not to create new models, but rather to test the utility of established computational models of B cells that have not previously been applied to lymphoma (10, 11). We performed no parameter fitting, as these parameters have been accumulated and validated across multiple cellular contexts and cell types (10, 12, 13, 33). We assume that parameters, other than those affected by mutations, remain consistent between healthy B cells and cancerous B cells. We also assumed that all mutations within patients have the same effect size, equivalent to one extra copy or loss of copy of each gene. The reality will be more complex, and we expect that identifying and quantifying the impact of each mutation on each gene would improve the utility of the model. Increasing the scope of the simulations, such that more recurrent mutations can be directly assigned to model parameters is also likely to improve the utility of this approach. However, the current model is clearly able to identify a population of very poor prognostic patients that would not have been identified using mutational clustering and COO alone.

Of note, the B cell receptor (BCR) and toll-like receptor (TLR) signalling pathways, although not explicitly modelled here, converge on NEMO-IKK and are the target of new therapeutic advances and recurrent mutations. Similarly, epigenetic regulators such as EZH2 are recurrently mutated but not explicitly modelled. Further work is also required to build on this model and simulate how treatment may perturb biological networks harbouring mutations. Such approaches may provide mechanistic insight into the development of relapsed DLBCL and provide insight into second-line treatments. Simulating the effect of targeted inhibitors in the patient-specific simulations presented here may provide insights into which patients will respond to alternatives to R-CHOP, however, limited clinical data is available to validate such predictions.

Performing large-scale computational simulations can be computationally challenging and require substantial computational resources given the size and complexity of the molecular networks simulated here (194 equations, and 563 parameters). Previous work has applied logical modelling (molecular components are discretised into high/medium/low) to B cell lymphoma to overcome this challenge (40). Here, we found a continuous modelling approach was able to identify many gene dose-dependent effects that could not be identified with logical modelling, including how chromosomal gain and amplification confer distinct prognoses. Despite the requirement for substantial computational calculation, we found that simulating just 6 hours of a single B cell per patient (simulating 100 cells required ∼30 minutes of real-word time on a 2.1 GHz CPU) enabled us to stratify patients. The ability to perform short simulations is enabled by experimental results that demonstrate that B cell molecular network states are rapidly established and reliably inherited across multiple generations of cell division (10). It is likely that clinical implementations of computational modelling could provide additional insight based on simulations without specialised hardware and without delaying treatment.

Beyond DLBCL and MM, the modelling methodology described here may have utility in other tumour types when mutational data and comprehensive, experimentally validated simulations are available. Many of the recurrent mutational events in DLBCL and MM are common cancer-associated mutations. cMyc deregulation is involved in the development of a wide variety of cancers (41), Bcl2 is dysregulated in many malignancies including breast cancer and gastric carcinoma (42, 43), and NF-κB is implicated in numerous cancers (44). A mathematical model, simulating 17 cancer types, estimated that the number of carcinogenic mutations (hits) varies from two to eight depending on the cancer type (45). With the increasing availability of cancer genome data and the evolution of computational methods to identify the mutational burden of cancer patients, the approach described here is rapidly becoming feasible in numerous cancers. We expect these approaches to be most impactful in mutationally heterogeneous cancers, such as breast cancer, where heterogeneity challenges early diagnosis, treatment selection and prognosis prediction (46).

The ultimate goal of computational systems biology approaches such as the one presented here is to enable personalised medicine by using models to get the right drugs to the right patients. Modern sequencing techniques provide an abundance of data, but we have yet to fully utilise it. Computational models with the power to translate patient-specific mutation data into personalised prognostic and treatment predictions may be required to enable truly personalised medicine approaches.

## Acknowledgements

The authors would like to thank Ulf Klein and Jana Wolf for critical discussions and Eleanor Jayawant for providing critical input into the manuscript. Funding for SM: Leukaemia UK John Goldman Fellowship (2020/JGF/003) and UKRI Future Leaders Fellowship (MR/T041889/1). Funding for AP: MRC Research Grant (MR/V009095/1). Funding for CP: Blood Cancer UK (23004).

## Author contributions

RN and SM performed computational analysis and modelling. RN, CP, AP and SM conceptualised the study, and EM, JJ and TC conceptualised the Fig. 3. AP, CP and SM supervised the project. All authors contributed to the article, critically revised the manuscript, and approved the submitted version.

## Data Availability

The datasets and computer code produced in this study are available in the following databases:

Modelling computer scripts: GitHub (https://github.com/SiFTW/norrisEtAl).

