## Supplementary File for "Patient-specific computational models predict prognosis in B cell lymphoma by quantifying pro-proliferative and anti-apoptotic signatures from genetic sequencing data"

**Supplementary Material**

#### Lay Summary of Methodology for Generating Computational Figures.

A computational model or simulation of a biological network consists of a text file containing computational code. This code can be run by specialist software, here we use a programming language called Julia. The code for a model consists of three parts: reactions, rate laws and tuneable parameters. Reactions define which biomolecules within the system interact, these can be visualised as the lines on the network diagrams (Fig 1A, 1D and S5). Rate laws define what mathematical function is used to simulate these interactions. These are predominantly basic mass action binding kinetics or Michaelis–Menten enzyme kinetics. The final piece of information needed are the kinetic parameters that define how fast each of these interactions proceed. Increasing specific parameters will increase the rate at which complexes bind or the rate at which genes are expressing.

The above information results in a model, defined by a number of equations: one for each biomolecule in the system. These biomolecules are predominantly proteins, protein complexes and mRNAs. There is an equation for each node in the diagrams in Fig 1A, 1D, and S5. These equations, called “differential equations”, describe the rate at which each biomolecule changes. Solving these equations generates time course trajectories predicting whether the concentration of each biomolecule increases or decreases over time. Each molecular interaction that impacts a biomolecule will add terms to its equation and therefore change the rate at which it either increases or decreases in concentration. Therefore, the arrows and interaction lines in the network diagrams (Fig 1A, 1D and S5) add to the equations of the nodes they interact with. In this way the molecular network diagrams are encoded in computer code that can be run to predict how concentrations will change over time.

**Fig. 1B** was generated by running the model shown in **Fig. 1A**. This model encodes just the apoptosis components from previous larger models [1]. The model was run with all parameters consistent with the published model except for BCL2 expression which was changed as shown in Fig. 1B. **Fig. 1C** was generated by running 1000 simulations of apoptosis that were all the same, except that the kinetic parameters were adjusted (randomly changed to be slightly higher or lower) for each cell. This parameter sampling captures cell to cell variability and results in different cells undergoing apoptosis at different times. This is shown as a distribution of death times. This was repeated for 1000 cells with published parameters (black), and 1000 cells with exactly the same parameters except for 1.5-fold increased BCL2 expression parameter (green). **Fig. 1E** was generated by running the model shown in **Fig. 1D**. This model encodes just the cell cycle components from previous larger models [1]. The model was run with all parameters consistent with the published model except for cMYC expression which was changed as shown in Fig. 1E. The top of Fig. 1E shows the result of running a simulation with published parameters for one whole cell cycle (solid lines). Also, on Fig. 1E is the result of running the same simulation but with cMYC expression increased by 1.5-fold. **Fig. 1F** was generated by running 1000 simulations of the cell cycle that were all the same, except that the kinetic parameters were slightly adjusted (randomly changed to be slightly higher or lower) for each cell. This parameter sampling captures cell to cell variability and results in different cell division times. This is shown as a distribution of division times. This was repeated for 1000 cells with published parameters (black), and 1000 cells with exactly the same parameters except for 1.5-fold increased cMYC expression parameter (green). For Fig. 1G, arrested cells were identified as cells that do not double in size prior to reaching M-phase.

**Fig. 2** was generated using complete model that contains the cell cycle, apoptosis, terminal B cell differentiation and NF-kB signalling [1]. A network diagram is shown in Fig S5 and summarised in 2A. The only change from the published model was that the NF-kB component was updated with a more detailed model of NF-kB signalling [2]. As this model contains both the cell cycle and apoptotic signalling networks, cells simulated this way can undergo apoptosis or mitosis. Apoptosis is triggered when cleaved PARP increases, and this is simulated by ending the simulation of that cell. Mitosis is triggered by CDH1 increasing at the end of M-phase and results in a daughter cell being simulated. The final concentrations of every biomolecule in a cell are used as the initial concentrations for the daughter cell simulation. This modelling approach is known as "agent-based" modelling. Simulations start with 100 cells with slightly randomised parameters (as described above for Fig. 1). Some cells will undergo apoptosis, while others will divide. The result is cell numbers changing over time as shown in **Fig. 2A**. In **Fig. 2B** (left) we ran a 100-cell simulation with published parameters (blue) and then with exactly the same parameters but cMYC expression increased 1.5-fold (gold), BCL2 expression increased 1.5-fold (red), and both increased simultaneously (green). We repeated this with a 5-fold increase (right). In **Fig. 2C**, we repeat the above, changing cMYC and BCL6 expression. In **Fig. 2E**, we repeat the above, changing MCL1 and CKS1B. As MCL1 is not explicitly included in the model, increased MCL1 was modelled as an increase in BCL2 expression. Similarly, CKS1B is not explicitly modelled but increased CKS1B is known to increase p27 degradation and was modelled in this way [3].

**Fig. 3A** was generated by running the model with published parameters (black dashed), with cMYC expression increased by 1.5-fold (orange), BCL2 expression increased by 1.5-fold (blue), and input IKK activity increased by 1.5-fold (green). Because IKK activity increased NF-kB activity, and both cMYC and BCL2 are NF-kB target genes, the simulation with increased IKK resembles a "double hit". **Fig. 3B** shows the approach we used to create personalised models. We used the whole exome sequencing dataset published by Chapuy *et al.* 2018. Mutations were downloaded from CBioPortal where the data was deposited. We only downloaded data for genes that impact genes/proteins within the model (list of genes downloaded: https://github.com/SiFTW/norrisEtAl/blob/main/geneList.txt). Each mutation was checked using a tool called OncoKB [4, 5], and only mutations annotated as "likely oncogenic" were included in subsequent analysis. Additional data quantifying copy number changes in chromosomal regions was downloaded from the supplement of the paper. The result of the above was a text file containing every (likely functionally significant) mutation and every copy number change in each patient. Each of these mutations was then mapped to model parameters. Some of these mappings were trivial, for example copy number gains in BCL2 map to the BCL2 expression rate parameters. Some mappings involve assigning mutations to the most biologically similar parameter, for example, we also map MCL1 copy number gains to the BCL2 parameter. The full mapping of mutations to model parameters is available on the GitHub repository (https://github.com/SiFTW/norrisEtAl/blob/main/muts2Params.csv). The result of this mapping was a parameter file for each patient. Every patient's parameter file was identical except for parameters impacted by mutations. The connections within the molecular network were assumed to be consistent across all patients, therefore the reactions and rate law definitions did not change between patients. The patient-specific parameters and patient-agnostic reactions and rate law definitions were combined to create patient-specific models.

In **Fig. 4A**, each patient-specific model was run and the concentration of anti-apoptotic proteins (renamed from Bcl2), and pro-proliferative proteins (renamed from cMYC) were plotted. Patients were stratified as being above or below the mean abundance of each of these factors at 6 hours. The survival of above average and below average patients is shown in **Fig. 4B**. Patients above average for both anti-apoptotic and pro-proliferative (AAPP) factors were identified in **Fig. 4C** and their clinical survival from Chapuy et al. is shown in **Fig. 4D**. The remaining Figs. use clinical data (downloaded from the cBioPortal data of Chapuy et al.) and mutational clusters (assigned by Chapuy et al.) in combination with the model-based stratification of patients as AAPP.

### Supplementary Modelling Methodology

**Model background.** The computational model used here consists of the agent-based ordinary differential equation (ODE) model of Roy, Mitchell et al. 2019 [1], which was converted from MATLAB to Julia to aid with reproducibility and accessibility without proprietary software licenses [6]. The NF-kB module of the model was replaced with an updated NF-kB model that includes non-canonical NF-kB signalling and additional inhibitors of NF-kB such as IkBδ/p100 [2]. As the inputs (NEMO:IKK activity) and outputs (nuclear RelA:p50 and cRel:p50) were consistent across the existing and newly introduced NF-kB module no additional reactions or parameters were required to update this model. The full description of the model parameters, reactions and rate laws is provided in the GitHub repository (https://github.com/SiFTW/norrisEtAl/), along with Jupyter notebooks to generate each Fig. The model's scope and reactions are depicted in Fig S5. This model was used to simulate individual cells in Fig. 1,3 and 4, as an agent-based model in Fig 2.

**Model construction.** To aid reproducibility the published MATLAB model, which contained reactions, discrete events, and parameters across multiple ".m" files, was converted into three CSV files for each module: reactions.csv, ratelaws.csv, and parameters.csv. These three CSV files were produced for the cell cycle, apoptosis, NF-kB, and differentiation modules. Individual modules were used in Fig 1. These modules were linked together using a further linking module to define the reactions reactions that contain components from multiple modules (e.g. cMYC being induced by NF-kB). The modules were combined by combining the CSV files to create three CSV files encoding the reactions, rateLaws, and parameters of the comprehensive model (Fig S5).

These CSV files are then converted to an executable model using bespoke Python code we have made available on Github (https://github.com/SiFTW/CSV2JuliaDiffEq). CSV2JuliaDiffEq uses the above CSV files to assemble a system of differential equations in Julia syntax compatible with DifferentialEquations.jl [7], and ensures model validity through guaranteeing mass conservation. The model is both constructed and solved within Jupyter notebook available on the Github repository.

**Apoptosis Modelling.** Apoptosis modelling in Fig. 1 was performed using just the apoptosis module (effectively isolating apoptotic signalling from the larger model). This model was simulated for $1x10^5$ minutes to reach equilibrium. Consistent with previous approaches, during this equilibrium phase, the concentration of TRAIL was set to 0 and BCL2 mRNA was set to 277*[*BCL2 modifier*] to ensure the cell did not undergo apoptosis during the equilibrium phase [1, 8]. [*BCL2 modifier*] was set to the values indicated in Fig. 1 (0.5, 1, 1.5 etc). The final concentrations from the equilibrium phase were used as initial conditions for the simulations shown in Fig. 1, except TRAIL was set to 1. Cell death was identified as the first time point at which over 10% of PARP was cleaved.

Simulations in Fig 1C were initiated with 1000 founds cells, with cell-to-cell variability created by sampling initial conditions as described previously [1, 8]**.**

**Cell Cycle Modelling.** Cell cycle modelling in Fig. 1 was performed using just the cell cycle module (effectively isolating the cell cycle from the larger model). As this model undergoes oscillatory behaviour it was simulated until a limit cycle was achieved or $1x10^6$ minutes, whichever came first. A limit cycle was defined when there was no disenable change in cell mass (within 3 decimal places) over 5 consecutive cell divisions. The final conditions of the limit-cycle phase were used as initial condition of the simulations shown in Fig. 1. During the pre-simulation phase a portion of cells did not double in size prior to division, and an event was added to detect this and arrest the cell cycle. The number of these cells was quantified in Fig. 1G. Cell cycle phase transitions were detected algorithmically using the maximum values of CDH1, CycA and CycB (full code available on the Github repository https://github.com/SiFTW/norrisEtAl/blob/main/Fig.1/CellCycle_WTvsMycOE.ipynb).

Simulations in Fig 1F were initiated with 1000 founds cells, with cell-to-cell variability created by sampling expression rate parameters as described previously [1, 8].

**Agent based modelling methodology.** The agent-based, mutli-scale modelling methodology was consistent with Roy, Mitchell et al. 2019 [1]. The code is available on the Github repository, and a Dockerfile is provided to ensure consistent versions of all software packages are used. Briefly each agent had two continuous callbacks representing cell division and cell death. The cell death callback was triggered at the first time point when cleaved PARP (cPARP) exceeded 2500. The cell cycle callback was triggered when CDH1 crossed 0.1 in the positive direction. The result of the cell division call back was the end of the simulation of that cell. The result of the cell cycle call back was to simulate cell division by stopping the simulation of the cell and introducing a daughter cell into the simulation. Here, we assumed perfect inheritance of all biomolecules across cell division and therefore the daughter cell's initial condition was copied from the final state of the mother cell. Due to this perfect inheritance only a single daughter cell was introduced representing both daughter cells. To track the number of cells represented by each daughter a cell each cell had a property assigned termed "cellsRepresented", which was calculated as $2^{(generation-1)}$. Simulations in Fig. 2 were initiated with 100 founds cells with cell-to-cell variability created by sampling expression rate parameters as described previously [1, 8]. As the model contains a constantly cycling cell cycle, that reaches a limit cycle rather than a steady state, an equilibrium phase was run with basal input (NEMO:IKK activity = 1%) for 1440 minutes prior to the start of the simulation. At timepoint zero IKK input is transiently increased using an input curve consistent with the published model.

**Patient simulations.** To create simulations of individual patients we created patient-specific parameters files using data from Chapuy et al. 2018. Data for 135 patients was available to download from cBioPortal. A gene list was created using all the recurrently mutated genes identified in the study, restricted to genes that could be mapped to model parameters. The gene list contained 64 genes and is available on the GitHub repository (https://github.com/SiFTW/norrisEtAl/blob/main/geneList.txt). The gene list was input into cBioPortal to select all patients who had a mutation affecting at least one of these genes (113 patients). The per-patient mutational data was downloaded as a tab-separated value file from cBioPortal. Each of these mutations was input into OncoKB, to determine whether it was annotated as "likely oncogenic". Only mutations meeting or exceeding this classification were included in subsequent analysis. Supplemental files from Chapuy et al. 2018 that described both focal and chromosomal arm-level copy number changes were also downloaded. For each gene in the gene list we identified whether it was affected by copy number changes due to these chromosome-level copy number changes. The mutations from cBioPortal and copy number changes from the paper's supplement were combined to give a comprehensive list of mutations for each patient. An additional file was created that described the mapping of each of these mutations to modelled parameters. This mapping assigned multiple mutations to the same model parameters. For example Myd88:L265P and CD79B:Y197D were both mapped to the parameters controlling NEMO:IKK activity. In this file all gain-of-function mutations (identified from the literature) were increased the parameter they were mapped to by 1.5-fold, while loss-of-function mutations reduced the parameter by 0.5-fold. When multiple mutations impacted the same model parameter in a single patient these effects were multiplied together. The above process resulted in 113 patient-specific parameter files that differed only by parameters impacted by mutations in the patient they represent. These were combined as described above to create patient-specific executable Julia models. These models were run for an equilibrium phase ($1x10^6$ minutes) prior to the plots shown in Fig. 4.

Supplementary Figures

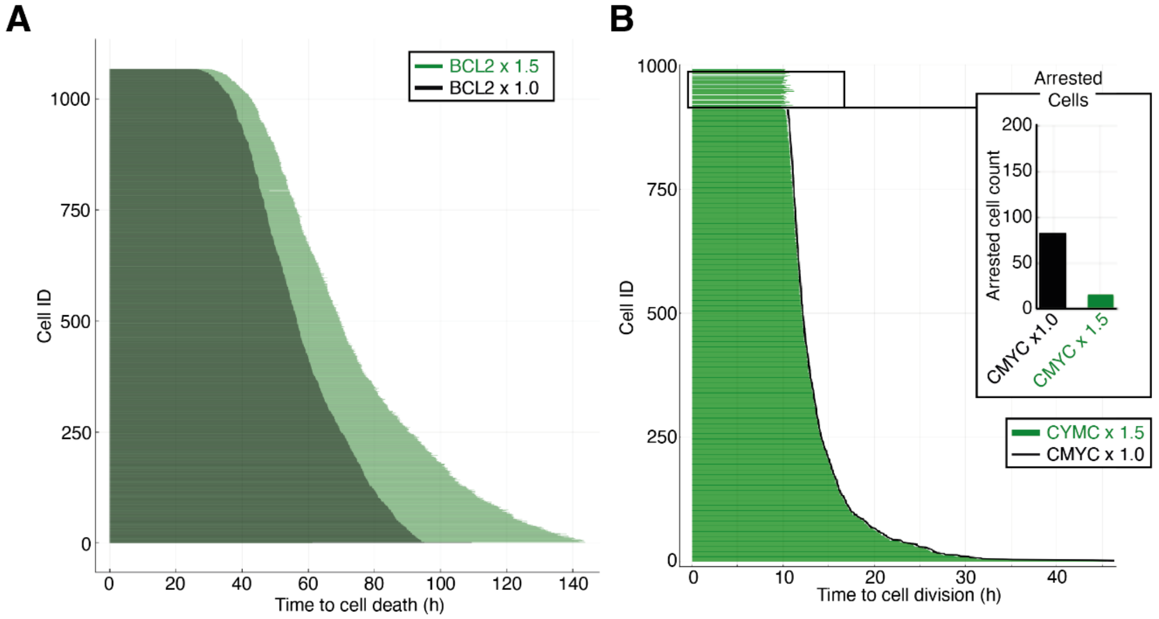

**Figure S1. Computational simulations of heterogeneous cell populations with over-expression of BCL2 and cMYC. A)** Simulations of a heterogeneous population of the apoptotic network (Fig. 1A) in individual B-cells with expression of BCL2 unchanged (1x black) or increased approximating one extra copy of the gene (1.5x green). Time to death was calculated by the value of cleaved PARP increasing as described previously [8]. **B)** Simulations of a heterogeneous population of the cell cycle network (Fig. 1D) in individual B-cells with expression of cMYC unchanged (1x black) or increased approximating one extra copy of the gene (1.5x green). Time to division was calculated by the value of CDH1 increasing as described previously [8]. Inset: number of cells in cell cycle arrest in each condition. Cell arrest was triggered by cell mass not increasing to twice the initial size prior to cell division.

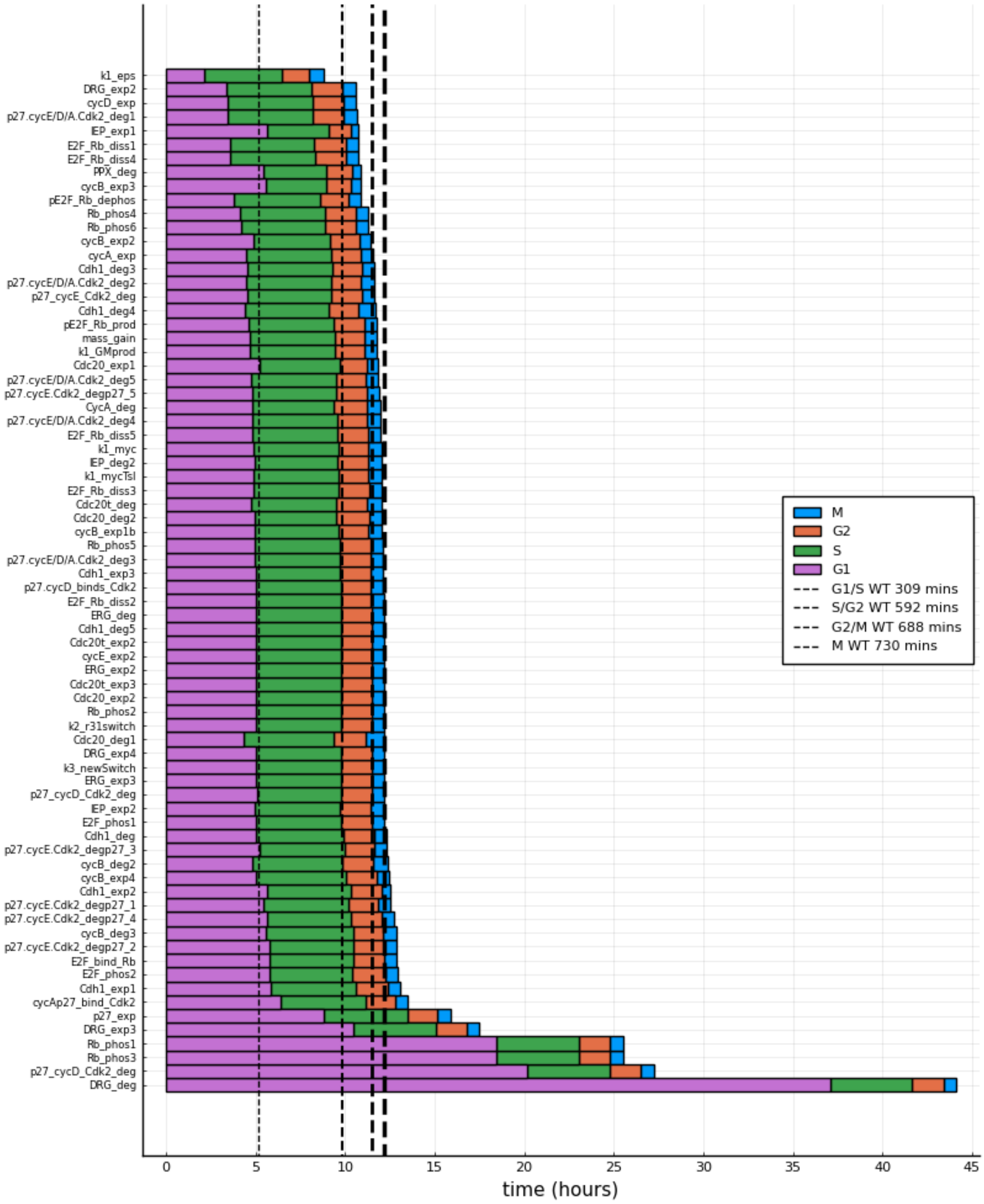

**Figure S2. Sensitivity analysis of the cell cycle model (Fig. 1D) predict that overexpression of cMYC speeds up the cell cycle but has a relatively small impact compared to other components.**

Each stacked bar represents a simulation of the cell cycle model with the indicated parameter altered (value x1.5). Simulations are plotted in order of total cell cycle time (top to bottom). Colours represent the different phases of the cell cycle: G1 = growth phase 1; S = synthesis; G2 = growth phase 2; M = mitosis. Black dotted lines indicate phase transition times for a wild type (WT) or unmutated cell with parameters unaltered. Individual parameter names are indicated and described in detail in https://github.com/SiFTW/norrisEtAl. Note the location of k1_mycTsl and K1_cMYC.

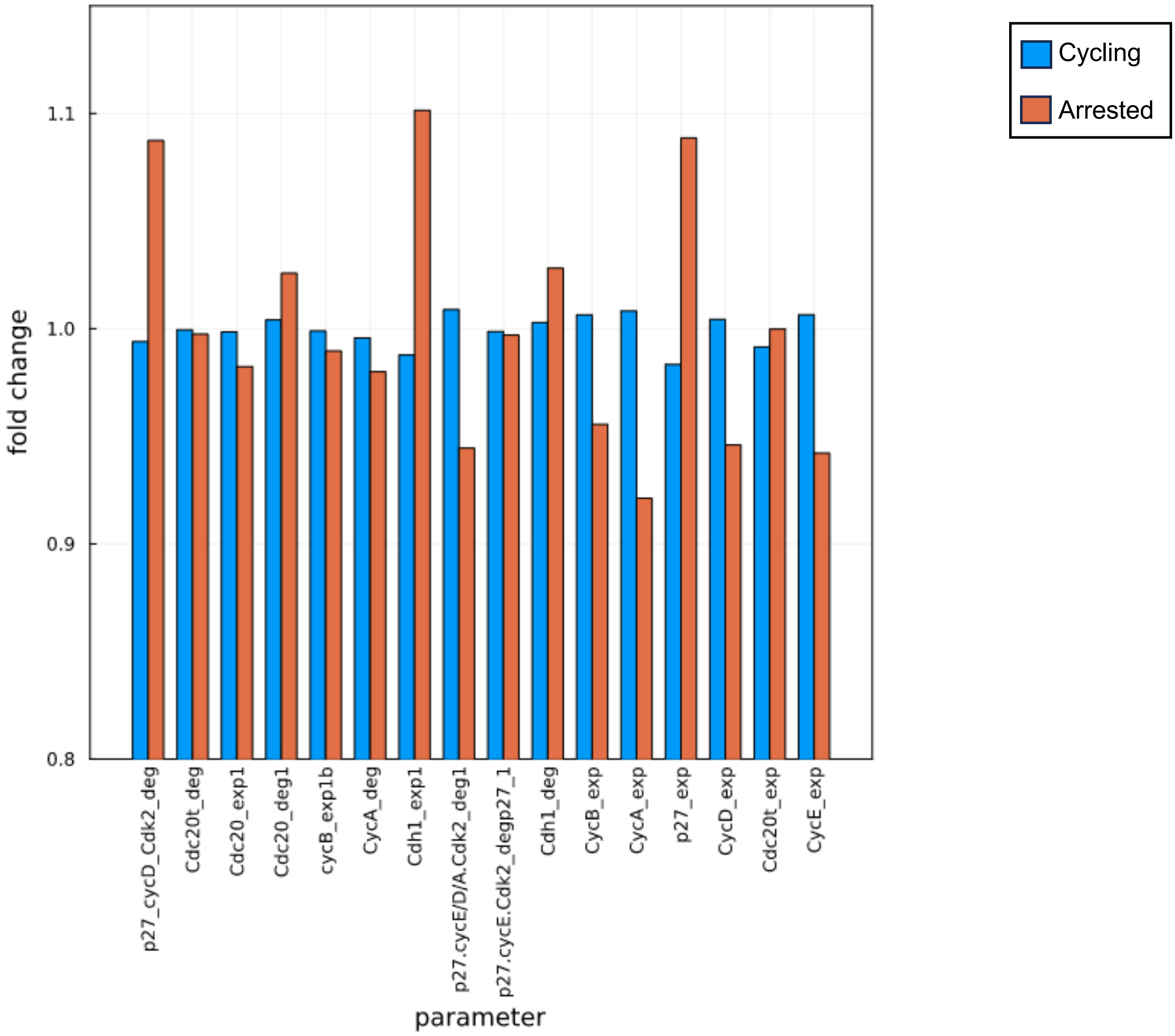

**Figure S3. Arrested cells have higher expression of p27 and Cdh1 than cycling cells in simulations of the cell cycle.**
Mean average values for all parameters distributed in simulated heterogeneous cell populations were calculated for arrested and cycling cells. These mean values were then compared to undistributed values and the difference plotted. Bars of fold change values for cycling/normal (blue) cells and arrested (orange) cells compared to undistributed values are plotted together along with a parameter names (x axis) for each parameter.

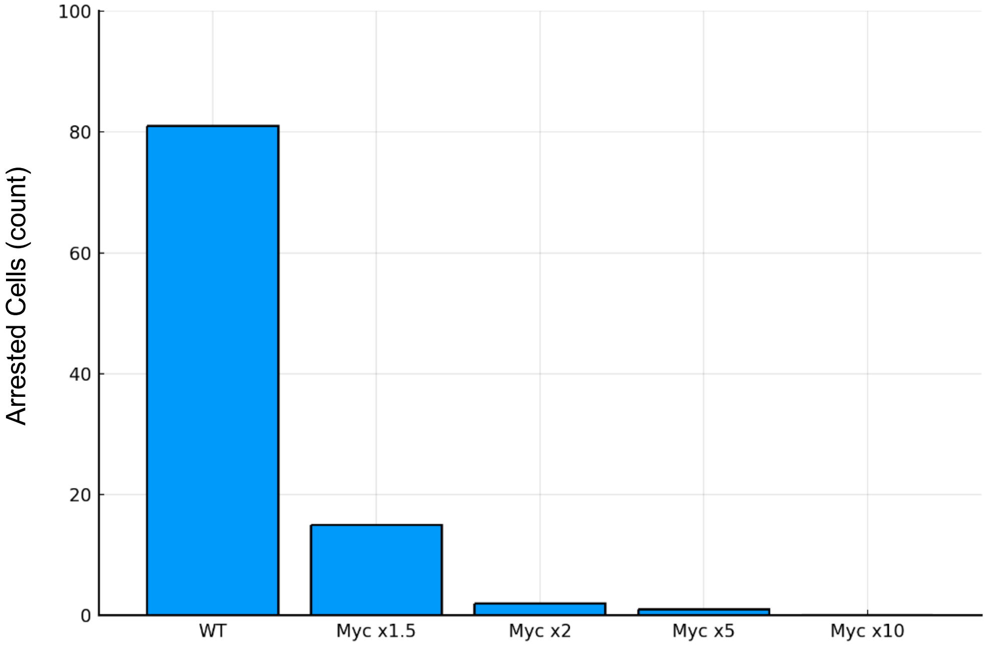

**Figure S4. Cell fate switching from arrested to cycling increases proportional to cMYC expression level**.
Bars indicate the number of cells in simulations of heterogeneous populations that have switched to an arrested state. The number of arrested cells is inversely correlated with cMYC expression level. At 10 times cMYC expression all cells that were arrested in the WT simulation are now cycling.

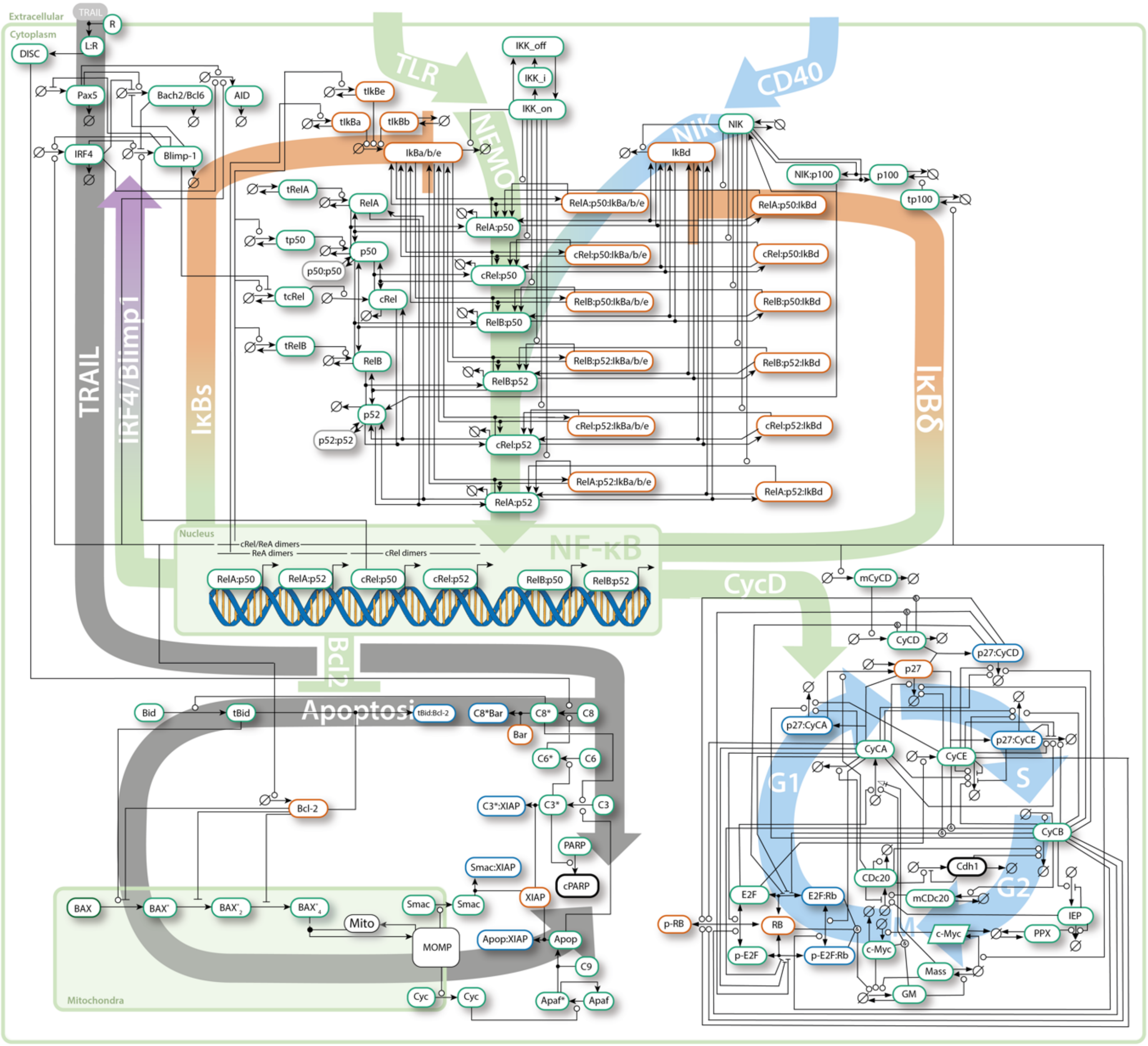

**Figure S5. Diagram of the scope of the multi-scale model of B-cell fates**.

All reactions, molecular species and parameters are maintained from previous publications [1]. The NF-κB component was updated from Roy et al. to include the more comprehensive reaction scheme from Tsui 2014 [9]. Some expression and degradation reactions are omitted for clarity. The full model is provided on Github https://github.com/SiFTW/norrisEtAl.

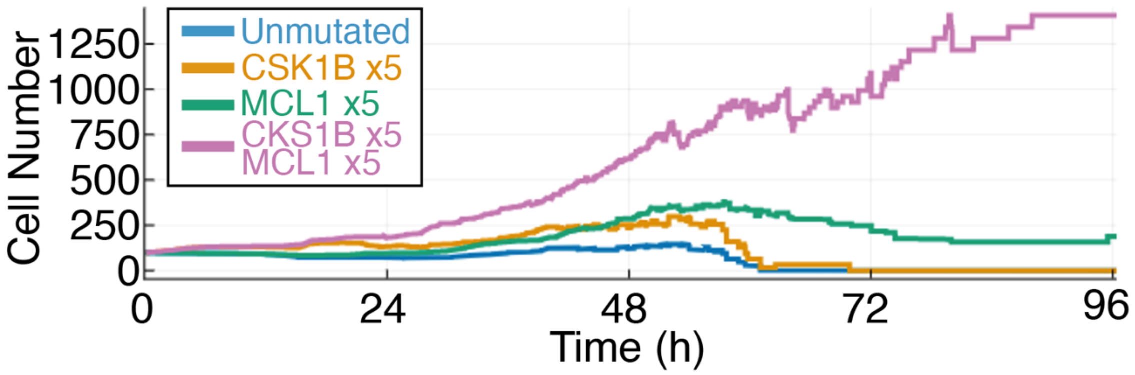

**Figure S6. Extended simulations of gain1q Multiple myeloma**.
Cell population size (cell count) over time for wild type (blue), individual mutations (orange and green) and double hits (purple) of the indicated genes. Note that after 2 48h of simulation the input curve has returned to basal levels and therefore continued proliferation is independent of NEMO-IKK activation (Figure 2A).

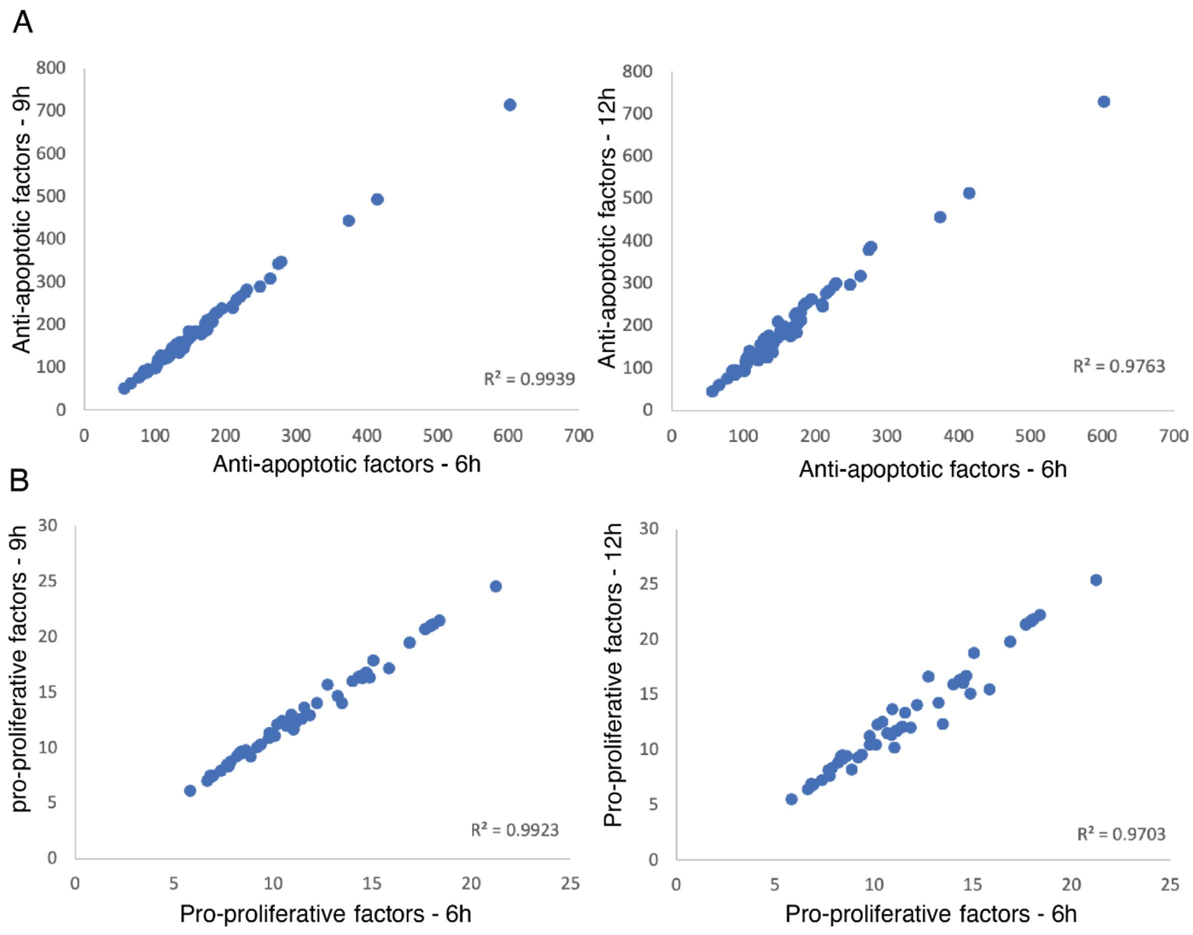

**Figure S7. Correlations between simulated concentrations of pro-proliferative and anti-apoptotic factors at 6, 9, and 12h.** Simulations were performed for 12h (Fig. 4A) and the concentration of the indicated molecular species selected at 6, 9 and 12h. Plotted is the correlation between these time points for all 113 patient-specific simulations. A) Correlation between anti-apoptotic factors. B) Correlation between pro-proliferative factors.

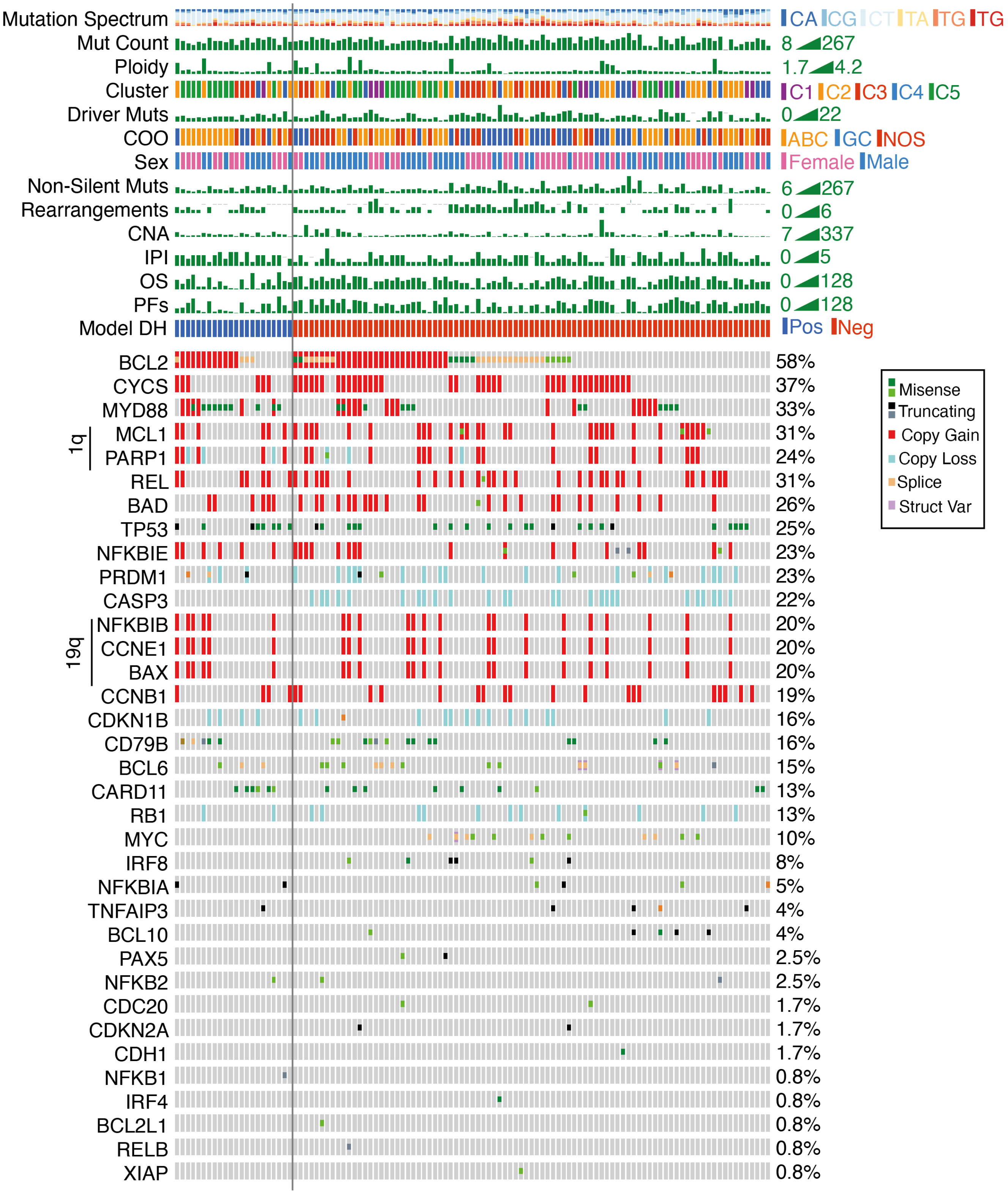

**Figure S8. Oncoprint of patients split by AAPP and "other" classifications (Fig. 4).** Patients on the left of the black line are anti-apoptotic and pro-proliferative (AAPP), while patients on the right are not.

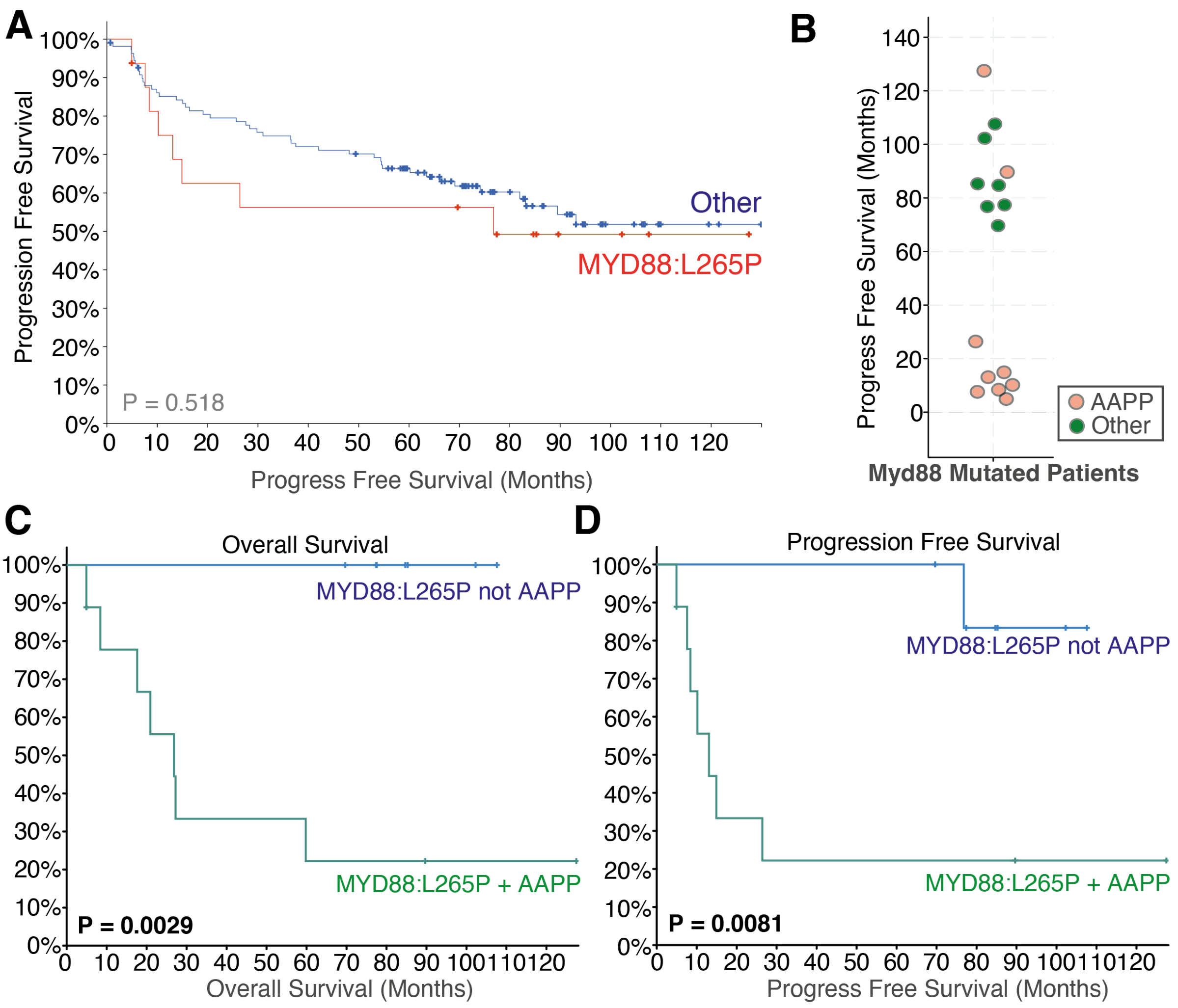

**Figure S9. Stratifying MYD88:L265P patients using computational modelling.** **A)** Kaplan-Meier plot showing Progression Free Survival (PFS) for patients with MYD88:L265P mutations and those without, patient mutation and outcome data from Chapuy et al. 2018 [10]. **B)** PFS for MYD88:L265P patients, showing bimodality of survival. MYD88:L265P patients who are also classified by the computational modelling as AAPP are shown in red, while MYD88:L265P patients not classified as AAPP are shown in green. **C and D)** Overall survival (C) and Progression free survival (D) for patients with MYD88:L265P mutations, stratified by whether computational modelling identifies them as AAPP (Green) or not AAPP (blue). P-value shown is from logrank test. AAPP = Anti-apoptotic and Pro-Proliferative.

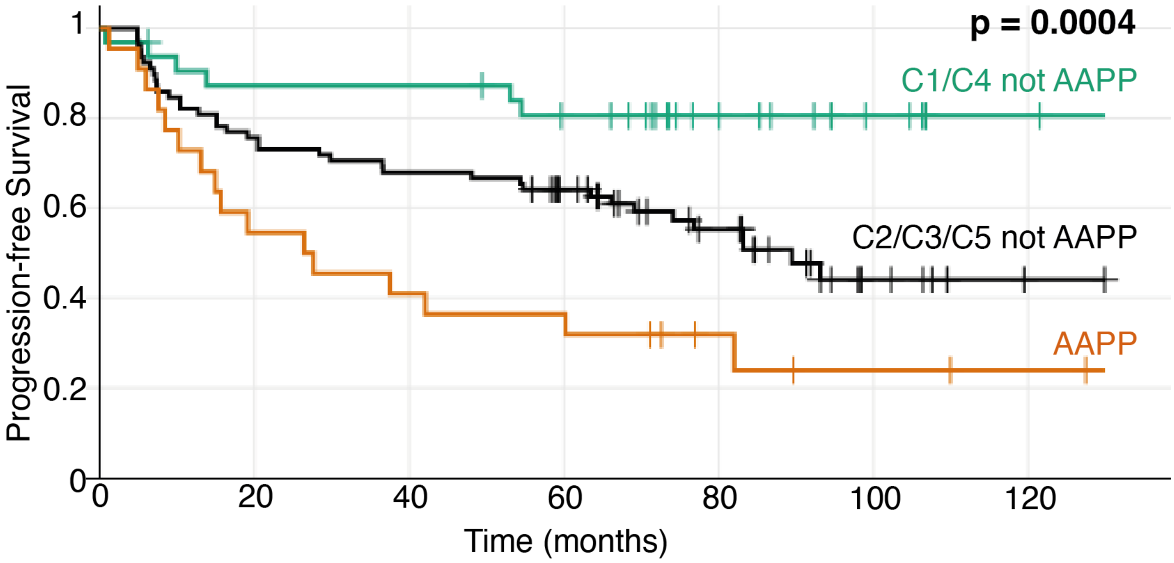

**Figure S10. Kaplan-Meyer plots of all patients stratified using a combination of computational modelling and mutational clusters.** This figure replicates analysis from Fig 5 but patients that cannot be modelled are included in their identified mutational clusters. Of the 135 patients available on cBioportal, 132 are modelled here as 3 lacked either overall survival data or mutational cluster.
